# Evolutionary tuning of a key helix drove androgen selectivity

**DOI:** 10.1101/2021.07.21.453223

**Authors:** C. Denise Okafor, Jennifer K. Colucci, Michael L. Cato, Sabab H. Khan, Kirsten Cottrill, David Hercules, Geeta N. Eick, Joseph W. Thornton, Eric A. Ortlund

## Abstract

The genetic and biophysical mechanisms by which new protein functions evolve are central concerns in evolutionary biology and molecular evolution. Despite much speculation, we know little about how protein function evolves. Here, we use ancestral proteins to trace the evolutionary history of ligand recognition in a sub-class of steroid receptors (SRs), an ancient family of ligand-regulated transcription factors that enable long-range cellular communication central to multicellular life. The most ancestral members of this family display promiscuous ligand binding due to their large ligand binding pockets, while more recently evolved SRs tend to have smaller cavities. Less obvious, however, are the forces driving the selectivity of highly similar ligands. A key example is the divergence between the progesterone and androgen receptors (PR, AR), which display a high degree of sequence similarity and yet display differential ligand preferences. This work uses the ancestral steroid receptor 2 (AncSR2), the common ancestor of all 3-ketosteroids and the ancestral androgen receptor 1 (AncAR1), the seminal androgen receptor, to explore the biophysical mechanisms that drove the evolution of androgen specificity. We determine that ligand specificity in androgen receptors is driven by changes in the conformational dynamics of the receptor as well as altered binding pocket interactions, with helix 10 (H10) playing a critical role in tuning ligand specificity.

## Introduction

The evolutionary history of a protein family is critical for deciphering the interplay between sequence, structure, and function in extant family members [1]. Ancestral sequence reconstruction (ASR) has permitted the resurrection and subsequent characterization of ancient proteins [2–5]. These studies have provided remarkable insight into the structural and biophysical mechanisms by which evolution drives functional divergence in paralogous proteins [6–9]. Studying protein function in extant proteins through mutational analysis is challenging; evolutionary neutral drift can often serve to ratchet protein function, leading to irreversible function switching [10, 11]. Investigation of the mechanisms driving a protein’s functional evolution reveal not only those residues responsible for the largest roles in function switching, but also the permissive mutations that allow the protein to access the conformational space to support the function-switching mutations [4, 6]. In an ancestral protein, these residue manipulations may be able to help “dial” a protein’s function, whereas these same changes in an extant protein may ablate function [12].

Steroid receptors (SRs) are a family of metazoan transcription factors that regulate a range of physiological processes in response to the binding of lipophilic cholesterol-derived hormones [14]. Ancestral Steroid Receptor 2 (AncSR2) represents the seminal 3-ketosteroid receptor, emerging around 450 million years ago (**Figure 1**) [13]. This receptor is promiscuously activated by a wide variety of lipophilic molecules, including all 3-ketosteroid hormones, though it displays a clear preference for 17-acetyl steroids (i.e. progestogens and corticosteroids) over 17-hydroxyl steroids (i.e. androgens) [15]. Thus, to specifically bind androgens, the androgen receptor (AR) evolved specific criteria to distinguish steroids based on C17 of their D-ring (**Figure 1**) [16]. To understand the molecular basis for discrimination of C17 substituents in the AR/PR clade, we leveraged an existing steroid receptor reconstruction [15] to resurrect the ligand binding domain (LBD) of the seminal androgen receptor, Ancestral Androgen Receptor 1 (AncAR1).

**Figure 1.**
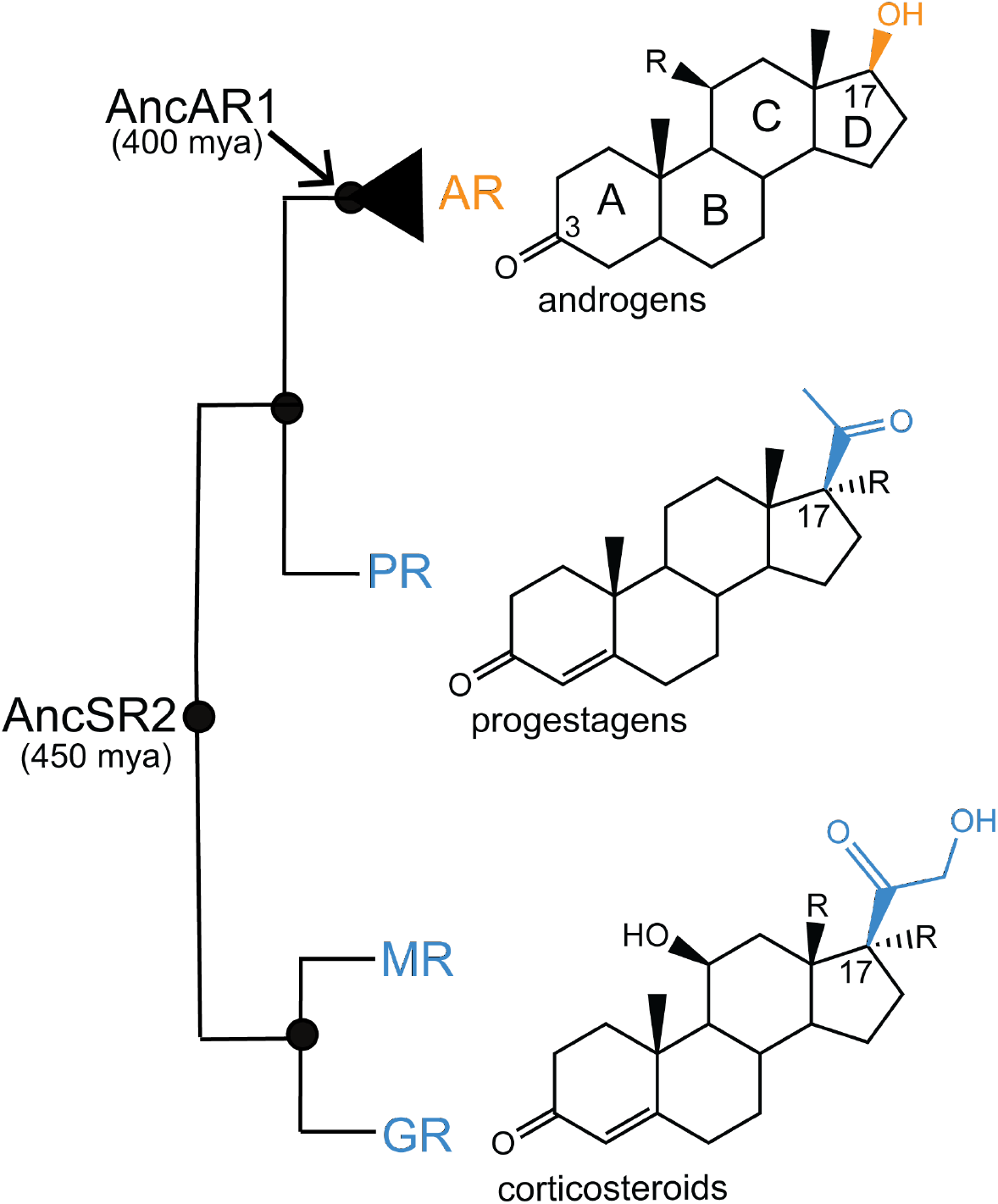
Evolution of androgen specificity. The phylogenetic relationship among 3-ketosteroid branch of steroid receptors showing the seminal 3-ketosteroid receptor AncSR2 and extant androgen, progesterone, mineralocorticoid, and glucocorticoid receptors. While the mineralocorticoid, glucocorticoid, and progesterone receptors are activated by 17-acetyl steroids corticosteroids and progestagens, the androgen receptor branch represents the emergence of 3-keto steroid receptors with specificity for C17-hydroxyl steroids (i.e. androgens). C17-acetyl groups are shown in blue, while C17-hydroxyl is shown in orange. The seminal androgen receptor AncAR1 is also indicated. For more comprehensive steroid receptor cladogram, see [13].

Here, we use structural, computational and in-cell assays to functionally characterize AncAR1, identifying and testing candidate historical substitutions to alter ligand specificity. Specifically, these analyses allow us to explore the historical mechanisms driving 3-ketosteroid receptor specificity. Our experiments demonstrate that a combination of historical mutations modulate receptor dynamics to partially recapitulate ancestral hormone preferences. Specifically, a leucine to threonine replacement in the LBP pocket eliminates steric bulk in the vicinity of the D-ring, allowing for 17-acetyl steroid binding. Additional substitutions on H10 alter helical dynamics, introduce salt bridges, and expand the binding pocket. Additionally, we leverage molecular dynamics (MD) simulations to identify evolutionary reverse substitutions which strengthen progesterone response relative to dihydrotestosterone (DHT). This work therefore highlights the ability to tune receptor function by exploiting evolutionary substitutions in a key helix.

## Results and Discussion

### Three historical substitutions restore progesterone activation in AncAR1

Consistent with its identity as the ancestor of all androgen-activated steroid receptors, AncAR1 is activated by the androgen DHT but not by progesterone, which differs only by its C17 substituent (**Figure 2A**). To understand how AncAR1 interacts with androgens, we determined the x-ray crystal structure of the AncAR1 ligand binding domain (LBD) in complex with DHT and a fragment of the TIF2 coregulator protein to 2.1 Å [17](**Figure 2B, S1**). The structure adopts the canonical “alpha helical sandwich” conformation observed for all steroid receptor LBDs. The C17 hydroxyl of DHT engages Thr208 and Asn36 in the ligand binding pocket (LBP) through hydrogen bonding. Both interactions are conserved in the human AR–DHT crystal structure [18] (**Figure 2C**).

**Figure 2.**
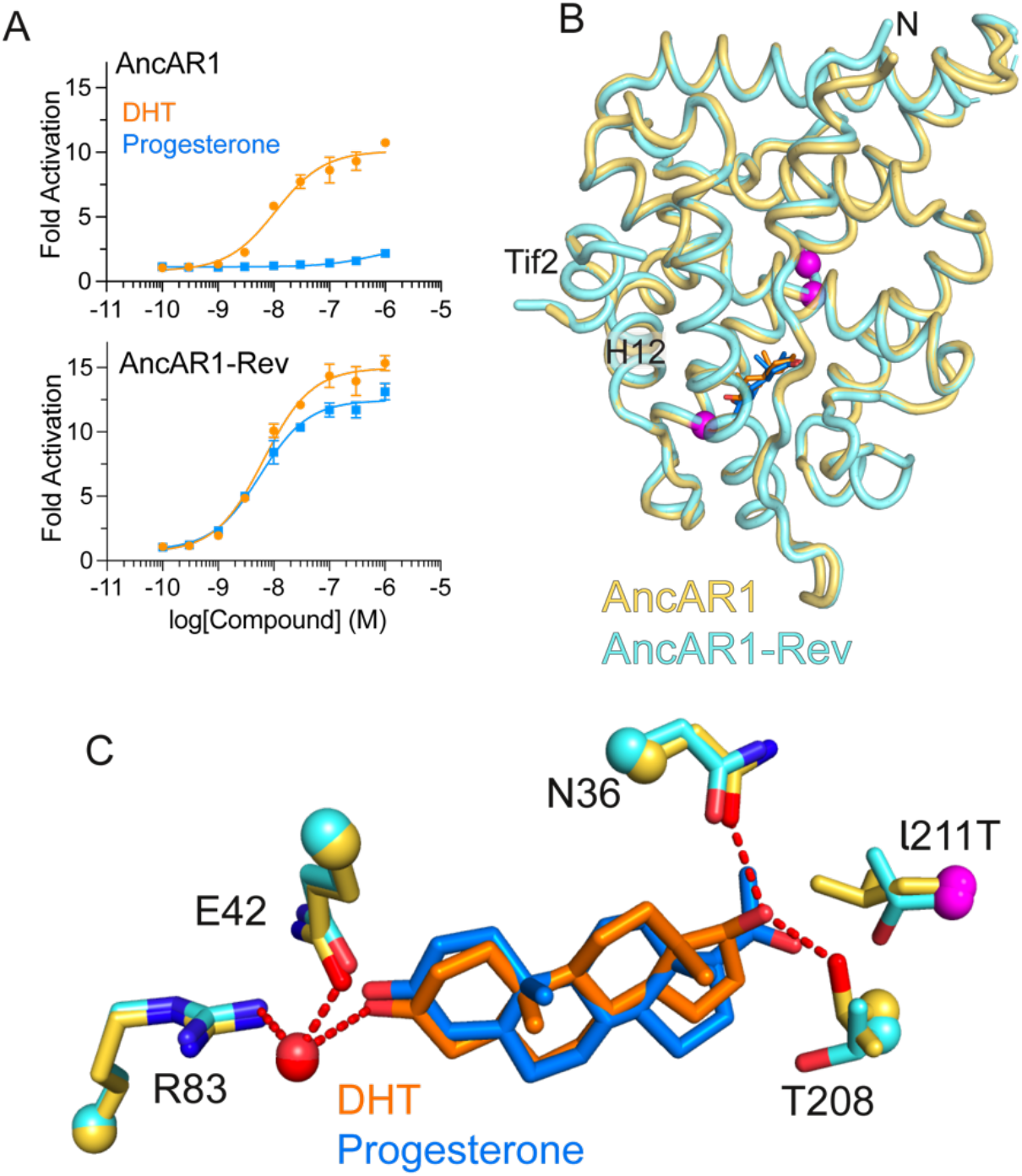
Progesterone activation is restored in AncAR1-Rev. A) AncAR1 is activated by DHT but not by progesterone in luciferase reporter assays. Data shown as mean ± SEM from two biological replicates. EC_50_ values are listed in Table 1. Evolutionary substitutions PRO199asp, ILE200leu, and LEU211thr (AncAR1-Rev) restore progesterone activation. Upper and lower case refer to derived and ancestral residues, respectively. B) X-ray crystal structures of AncAR1 and AncAR1-Rev LBDs are superimposed with substituted positions (199, 200, 211) highlighted in magenta. C) Residues that contact the steroid D-ring. DHT makes polar contacts (red dashes) with THR208 and ASN36 in the binding pocket, which are absent in progesterone. LEU211 in AncAR1 is replaced by thr211 in the AncAR1-Rev-progesterone complex.

To identify candidate historical substitutions that would reintroduce progesterone sensitivity into AncAR1, we performed a combined structure and sequence analysis across the steroid receptor family. We hypothesized that position 211, a leucine in all 3-ketosteroid receptors, but a threonine in AncAR1, would be a significant evolutionary mutation given its proximity to the D-ring of the ligand (**Figure 2C**). When introduced to AncAR1, the LEU211thr substitution slightly enhanced progesterone activation in a luciferase assay but also decreased DHT response (**Table 1**). Therefore, we tested the LEU211thr substitution in combination with other historical substitutions. This narrowed the set to three positions that restored full progesterone response to AncAR1: PRO199asp, ILE200leu and LEU211thr, referred to here as AncAR1-Rev (**Figure 2A**).

**Table 1.** EC_50_ values calculated from luciferase reporter assays to quantify DHT and progesterone activation of AncAR1 variants. 95% confidence intervals shown in brackets.

| Receptor | DHT |  | Progesterone |  |
| --- | --- | --- | --- | --- |
|  | EC<sub>50</sub> (nM) | E<sub>max</sub> | EC<sub>50</sub> (nM) | E<sub>max</sub> |
| AncAR1 | 11 [7.3, 16] | 10 | 590 [190, 4300] | 2.6 |
| AncAR1 + L211T | 48 [23, 100] | 10 | 120 [63, 240] | 6.6 |
| AncAR1 + P199D | 5.1 [3.9, 6.7] | 13 | 67 [34, 140] | 2.6 |
| AncAR1 + I200L | 4.8 [3.8, 6.1] | 11 | 50 [32, 82] | 6.3 |
| AncAR1 + L211T+P199D | 15 [8.0, 30] | 9.9 | 37 [19, 70] | 5.8 |
| AncAR1 + L211T+I200L | 16 [12, 21] | 11 | 12[9.8, 15] | 9.1 |
| AncAR1 + I200L+P199D | 2.3 [1.5, 3.6] | 14 | 20 [14, 29] | 8.7 |
| AncAR1-Rev | 6.5 [4.9, 8.7] | 15 | 5.7 [4.3, 7.6] | 13 |

To enable a detailed structural comparison between the ligand binding domains of AncAR1 and AncAR1-Rev, we determined the structure for AncAR1-Rev LBD in complex with progesterone to 1.5 Å (**Figure S1**). The two structures superimpose with a carbon alpha RMSD of 0.31 Å (Figure 2B). ProSMART analysis reveals a small degree of variability at the N-terminus (H1), the H8-H9 loop and on H12 (**Figure S2**). A-ring interactions, driven by a conserved network of H-bonds between the 3-keto oxygen, a conserved water, GLN42 and ARG83, are identical between both structures. The reverse substitution LEU211thr confers AncSR2-like recognition of progesterone by creating additional space to accommodate the larger C17 acetyl group. The critical polar interactions between DHT and ASN36 and T211 are not present in AncAR1-Rev-progesterone.

### Structural mechanisms underlying effects of substitutions

To determine the mechanisms by which the reverse substitutions restore progesterone activation, we compared the structures of AncAR1 and AncAR1-Rev. Two of the three substitutions, ILE200leu and PRO199asp, lie in the middle of H10 and are positioned to interact with the adjacent H7 **(Figure 3A-B)**. The ILE200leu substitution alters hydrophobic packing between the H10 and surrounding helices (**Figure 3A**), increasing the buried surface area as measured using PDBePISA (**Figure S2**). Because of their proximity to the ligand binding pocket, this change in packing may serve to stabilize an active conformation of AncAR1. Consistent with this proposed effect, the introduction of the single ILE200leu mutation into AncAR1 increases both maximum activation and potency of progesterone but has no effect on DHT response (**Figure 3C**). Additionally, accelerated MD simulations comparing ILE200leu AncAR1 mutant complexes to wildtype AncAR1 complexes show that conformational dynamics are altered in progesterone-bound receptors, but there are no differences between dynamics of DHT-bound wildtype and ILE200leu AncAR1 complexes (**Figure S3**).

**Figure 3.**
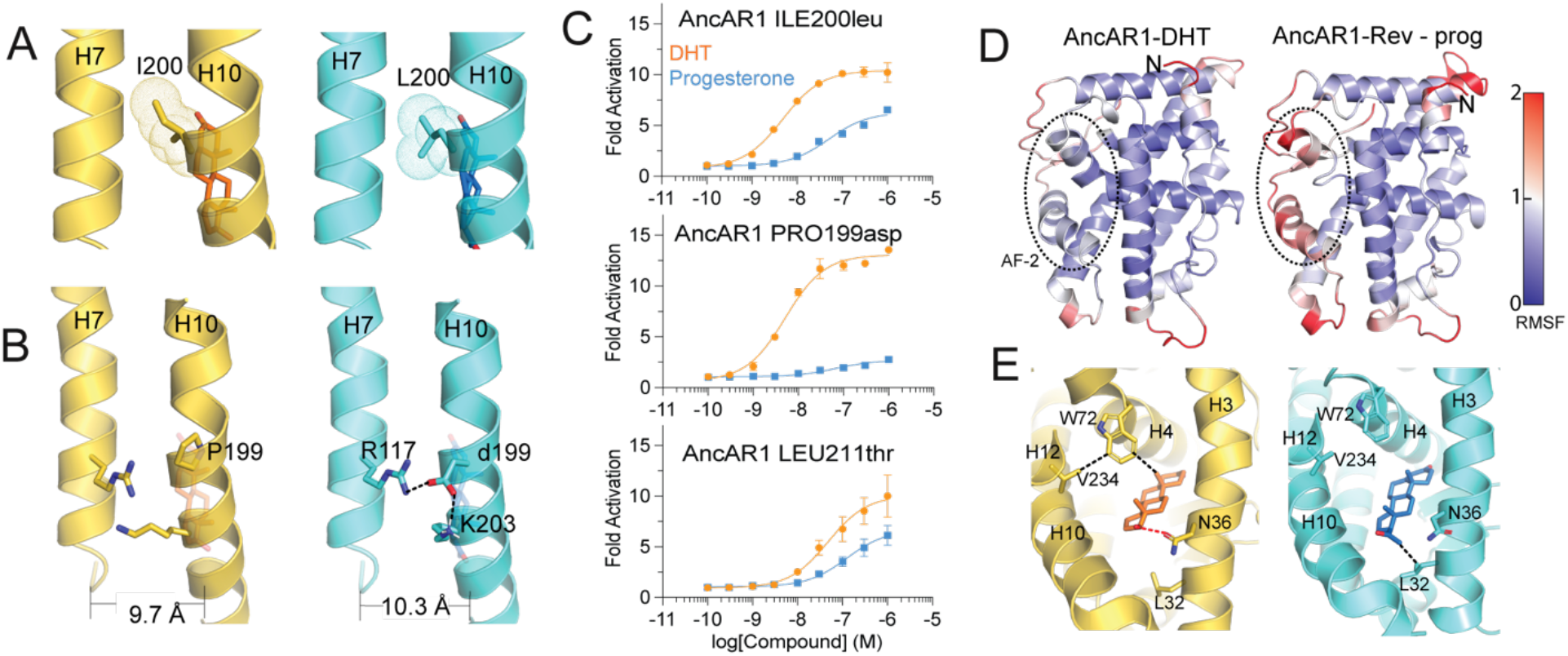
Reverse evolutionary substitutions stabilize AncAR1. A) ILE200leu substitution alters the helical packing between H7 and H10. B) PRO199asp substitution introduces two salt bridges and increases the H7-H10 interhelical distance. C) Luciferase reporter assays show that ILE200leu increases sensitivity to progesterone without affecting DHT-mediated activation. PRO199asp appears neutral relative to AncAR1. LEU211thr reduces DHT activation while enhancing progesterone response relative to AncAR1. Data shown as mean ± SEM from two biological replicates. EC_50_ values are listed in Table 1. D) Root mean square fluctuations analysis reveals that AncAR1-Progesterone displays increased fluctuations at the AF-2 surface E) Contacts analysis from MD simulations reveals that the instability results from the inability of progesterone to interact stably with TRP72, contributing to destabilizing H12.

The PRO199asp substitution eliminates the helix-breaking proline and introduces two salt bridges with ARG117 and K203 on H7 and H10, respectively (**Figure 3B**). These combined effects influence conformational dynamics in H10. Our accelerated MD simulations confirm that the introduction of the single PRO199asp substitution in AncAR1 alters conformational dynamics of both DHT- and progesterone-bound AncAR1 (**Figure S3**). While mechanisms are poorly understood, the vital role of conformational dynamics in driving the evolution of protein function is well established [19].

Both the crystal structures (**Figure 3D**) and MD simulations (**Figure S4**) show an increase in the H10-H7 interhelical distance. An increased H10-H7 distance increases the size of the pocket, enabling AncAR1 to support the larger progesterone hormone. Despite predicted dynamical changes in H10, the single PRO199asp mutation in AncAR1 is neutral, having virtually no effect on DHT or progesterone activation (**Figure 3C**). However, an epistatic effect is observed when PRO199asp is combined with either of the other two substitutions. PRO199asp+ILE200leu drastically increases activation by both hormones, while PRO199asp+LEU211thr only enhances response to progesterone (**Figure S4, Table 1**).

In addition to altering D-ring interactions within the binding pocket, the single LEU211thr is unique as the only individual substitution that weakens DHT response (**Figure 3C**). This reduction in DHT activation is mitigated when LEU211thr is combined with either of the PRO199asp or ILE200leu substitutions (**Table 1**), supporting the hypothesis that these residues stabilize an active conformation of AncAR1.

The presence of progesterone response even in LEU211 AncAR1 variants (i.e. PRO199asp, ILE200leu and PRO199asp/ILE200leu substitutions) suggests that the bulky leucine sidechain does not fully occlude progesterone binding. This observation implies that other mechanisms in play prevent progesterone activation in AncAR1. To further explore these mechanisms, we performed MD simulations on our AncAR1-DHT structure and an AncAR1-progesterone complex generated *via* modeling. A root mean square fluctuations (RMSF) analysis reveals increased dynamics at the Activation Function 2 (AF-2) surface of AncAR1-progesterone, a critical region of the receptor responsible for interaction with coregulators and downstream activity (**Figure 3D**). We used a contact analysis of the binding pocket to uncover a structural explanation for these dynamic effects. ‘Contacts’ are defined between two non-neighboring residues if their heavy atoms reside within 4.5 angstroms of each other for 75% of the MD trajectory. DHT engages ASN36 in a hydrogen bond that persists throughout a microsecond-long trajectory (**Figure 3E**). Progesterone does not form this hydrogen bond, instead engaging LEU32 in a van der Waal interaction via its bulky acetyl D-ring substituent. This interaction affects ligand engagement with TRP72 on helix 3 for both hormones, as DHT but not progesterone forms a contact with TRP72 in simulations. Widely conserved in nuclear receptors and crucial for its role in stabilizing H12, TRP72 is positioned to stabilize DHT in the pocket but unable to stabilize progesterone [20]. The loss of this stabilization in the AncAR1-progesterone complex therefore likely contributes to increased dynamics at H12 as observed in MD simulations.

### Forward trajectory is epistatically restricted

Having established roles for each of the three residues in the functional reversal, we investigated the reversibility of this evolutionary trajectory by determining whether the ‘forward’ substitutions in the AncSR2 background would abolish progesterone activation. We inserted the three substitutions (leu200ILE, asp199PRO, thr211LEU) into AncSR2 individually and evaluated transcriptional activation by DHT and progesterone (**Figure 4**). The leu200ILE substitution appears to be neutral, with no effect on DHT or progesterone response in AncSR2 (**Figure 4 A,B**). The asp199PRO substitution noticeably perturbs AncSR2 function, drastically lowering maximum activation for both ligands (**Figure 4C**).

**Figure 4.**
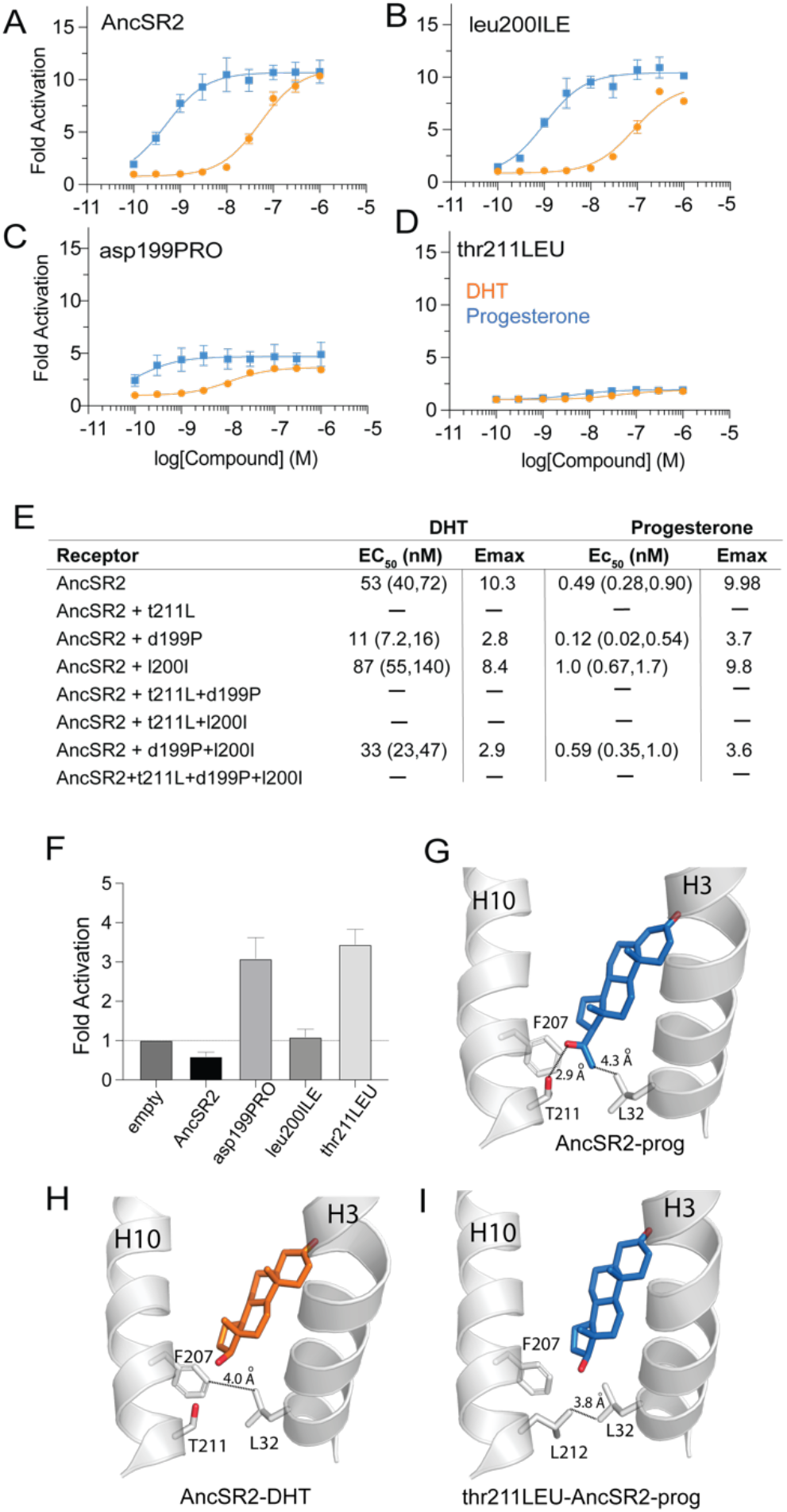
Effects of forward substitutions on AncSR2 activity. A) AncSR2 is activated by both progesterone and DHT. B) the leu200ILE AncSR2 mutant appears neutral with respect to progesterone and DHT activation. C) asp199PRO mutant drastically lowers maximum activation. D) thr211LEU mutant reveals no ligand-dependent activity. Data shown as mean ± SEM from two biological replicates. E) Activation data obtained from AncSR2 mutants in luciferase reporter assays. F) In the absence of ligand, both asp199PRO and thr210LEU AncSR2 mutants display constitutive activity. Data shown as mean ± SEM from two biological replicates. G) In AncSR2, progesterone interacts with residues on helices 3 (L32) and 10 (T212) to stabilize the pocket. H) In AncSR2-DHT, the pocket is stabilized by a H3-H10 interaction between F207 (H10) and L32 (H3). I) The thr211LEU substitution in AncSR2 allows direct H3-H10 contact without altering the size of the pocket, conferring ligand-independent activity to AncSR2.

The thr211LEU produces the strongest effect, conferring constitutive activity to AncSR2 (**Figure 4 E,F**). A structural inspection, along with analysis of our MD trajectories, suggests a plausible explanation for this effect: introducing leucine in the AncSR2 LBP forms a network of interactions that keeps the receptor in an activated state. leu32 on H3 and thr212 act as ligand sensors in the AncSR2 pocket; both can interact with the D-ring of progesterone (**Figure 4G**) but not DHT (**Figure 4H**). This network of interactions links H3 and H10 with the bound hormone in progesterone complexes. AncSR2-DHT complexes are stabilized instead by an interaction between leu32 and phe207 (**Figure 4G**). In the absence of ligand, the pocket collapses, evidenced by smaller volume observed in MD simulations of apo AncSR2 (**Figure S5**). With the thr211LEU substitution (**Figure 4I**), the bulkier leucine introduces a direct contact between leu212-leu32, stabilizing the H3-H10 interaction independently of ligand, while maintaining the ligand cavity (**Figure S5**).

We evaluated the effect of combined forward substitutions on transcriptional activity. All asp199PRO variants display a drastically reduced maximum activation, while all thr211LEU variants, including the triple-substituted AncSR2 are constitutively active (**Figure 4**). These observations indicate that additional epistatic substitutions must have preceded both thr211LEU and asp199PRO to ensure that functional switching could be tolerated in AncAR1 evolution.

### ASN36 is a critical residue that enables stabilization of productive complexes

Allosteric regulation is a key aspect of steroid receptor function. Upon hormone binding, transcription is mediated by ligand-induced conformational changes that lead to the recruitment of coregulator proteins to the AF-2. To enhance our understanding of how reverse evolutionary substitutions restore progesterone activation to AncAR1, we sought to determine how the sequence changes affect allosteric signaling between the ligand and AF-2 surface in the receptors. To predict and quantify the strength of allosteric signaling paths, we employed a suboptimal paths analysis of MD trajectories which uses calculated covariance between residue pairs to predict amino acids involved in signaling. Suboptimal paths in the AncAR1–progesterone complex are severely restricted compared with AncAR1-DHT (**Figure 5**). Closer analysis reveals the origin of these restrictions as a network of crucial edges between ASN36 and surrounding residues that are weakened in the AncAR1–progesterone binding pocket (**Figure 5B**). Weak ligand-ASN36 communication in AncAR1-progesterone propagates to H12/AF-2 via weakened edges between H12 and H3 residues ASN36 (**Figure 5C**) and GLU40 (**Figure 5D**). ASN36 forms a hydrogen bond with DHT persisting for 93% of the simulation compared to 0.03% in progesterone (**Figure 5E**). Conserved across all 3-ketosteroid receptors, ASN36 is part of a critical H-bond network that stabilizes steroids and H12 and is crucial for steroid receptor activation [21–24]. We propose that the lack of stable N36-ligand hydrogen bond formation in AncAR1-progesterone plays a role in impairing LBP dynamics and restricts signaling from the pocket. Consistent with restricted signaling, suboptimal paths are weaker in AncAR1-progesterone compared to AncAR1-DHT (**Figure 5F**).

**Figure 5.**
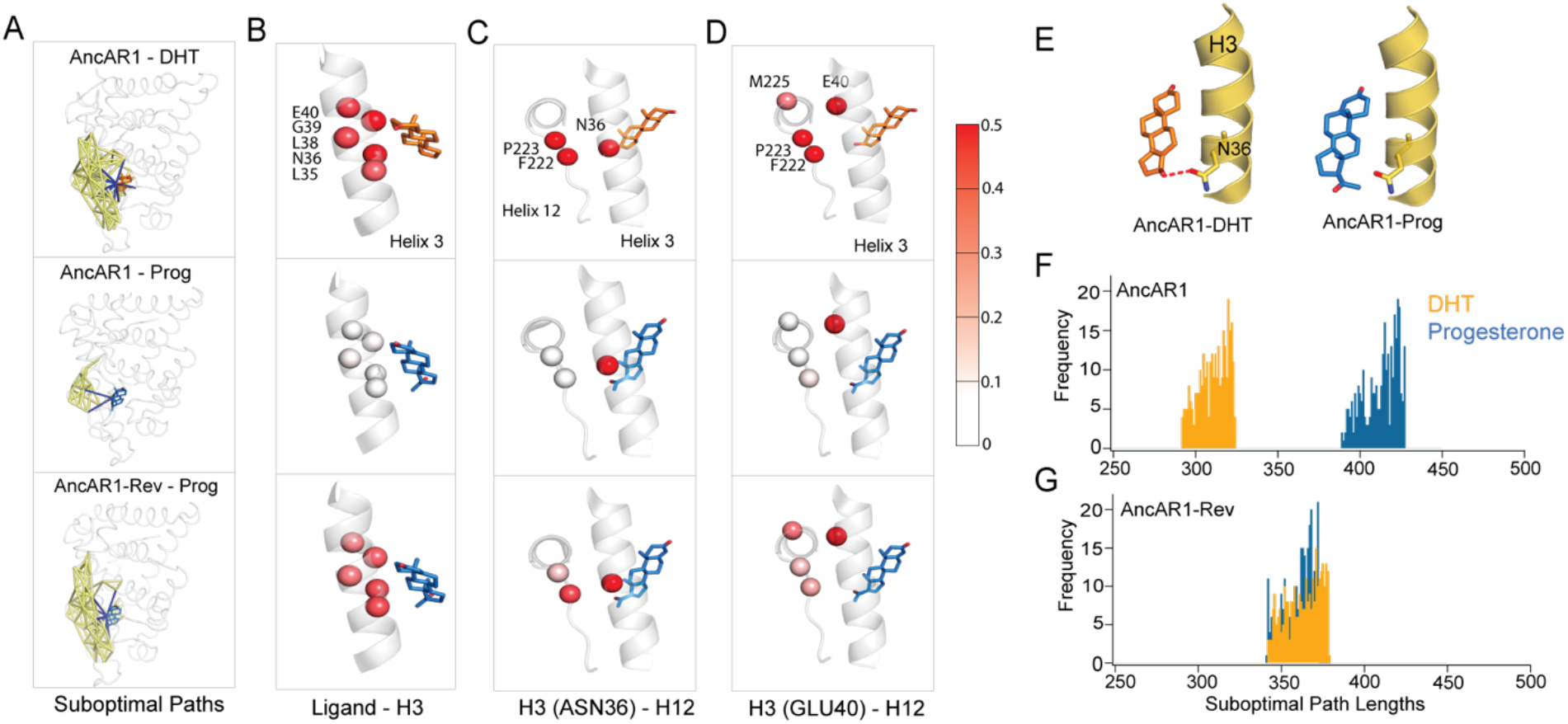
Allosteric signaling in AncAR1-progesterone is restricted. A) Suboptimal paths analysis derived from MD simulations are used to visualize allosteric communication between the ligand and AF-2 surface. Edges directly contacting ligand are shown in blue, all other path edges are yellow. Stronger communication (allostery) is indicated by a greater number of paths. Restricted signaling is predicted in the AncAR1-progesterone complex (middle), compared to AncAR1-DHT (top). Reverse substitutions restore enhanced communication in AncAR1-Rev progesterone complex (bottom). B) Strength of first-shell ligand interactions between the hormone and helix 3. Helix residues are colored by covariance between ligand and each residue. Covariance, calculated from MD trajectories, is greatly reduced in AncAR1-progesterone complex, but enhanced in AncAR1 rev complex. C-D. Strength of second-shell interactions between helices 3 and 12. Residues are colored by covariance between C) ASN36 and H12 and D) GLU40 and H12 residues. Similar to first shell interactions in B, signaling is also weakened between H3 and H12 in AncAR1-progesterone complex. This effect is reversed upon insertion of reverse substitutions. E) Simulations show that ASN36 of AncAR1 forms a hydrogen bond with DHT that persists for 93% of the simulation. Hydrogen bonding is not observed with progesterone. F-G) Histogram showing the shortest 1000 suboptimal paths between the hormone and helix 12 in AncAR1 and AncAR1-Rev. In AncAR1, path lengths are stronger in DHT than progesterone. In AncAR1-Rev, path lengths are equal for both complexes.

AncAR1-Rev enhances interactions between N36 and progesterone, permitting stronger signaling to AF-2 (**Figure 5E**). The edges are stronger, implying that covariance between these residue pairs is increased in AncAR1-rev-progesterone complex. The strength of suboptimal paths is equal for both AncAR1-Rev-Progesterone and AncAR1-Rev-DHT (**Figure 5G**), consistent with nearly equal transactivation potency of both hormones (**Figure 1B**). Simulations do not predict an increase in hydrogen bonding between progesterone and N36 (**Figure 5E**), suggesting that the enhanced correlation between progesterone and pocket residues is enabled, in part, by the increased stability of progesterone in the pocket resulting from the combined effects the reverse evolutionary substitutions.

### Two additional H10 substitutions restore ancestral selectivity

While AncAR1-Rev is strongly activated by both hormones, AncSR2 displays a clear preference for progesterone with ~ 100-fold lower EC_50_ than DHT (**Figure 1B, Figure 4A**). We sought to identify additional evolutionary substitutions to recapitulate the AncSR2 preference in AncAR1 (i.e. increasing progesterone activation relative to DHT). We analyzed inter-residue contacts in our MD simulations and identified four residues that made distinct contacts in DHT vs progesterone complexes (**Figure S6**). Thus, we hypothesized that one or more of these four positions: ARG202, LYS203, HIS205, and THR208 would modulate ligand selectivity (**Figure 6A**). We note that two of these four positions, HIS205 and THR208 correspond to prostate cancer mutation hotspots in human AR: residue numbers HIS874 and T877 respectively [25, 26]. To predict the impact of ancestral reversions of these four positions, we introduced the historical substitutions ARG202gly, LYS203gly, HIS205leu and THR208cys in the background of AncAR1-Rev and performed MD simulations of the receptors complexed with both DHT and progesterone. Suboptimal paths analysis revealed two states (ARG201gly and the HIS205leu/THR208cys double mutant) with stronger progesterone signaling (**Figure 6B**).

**Figure 6.**
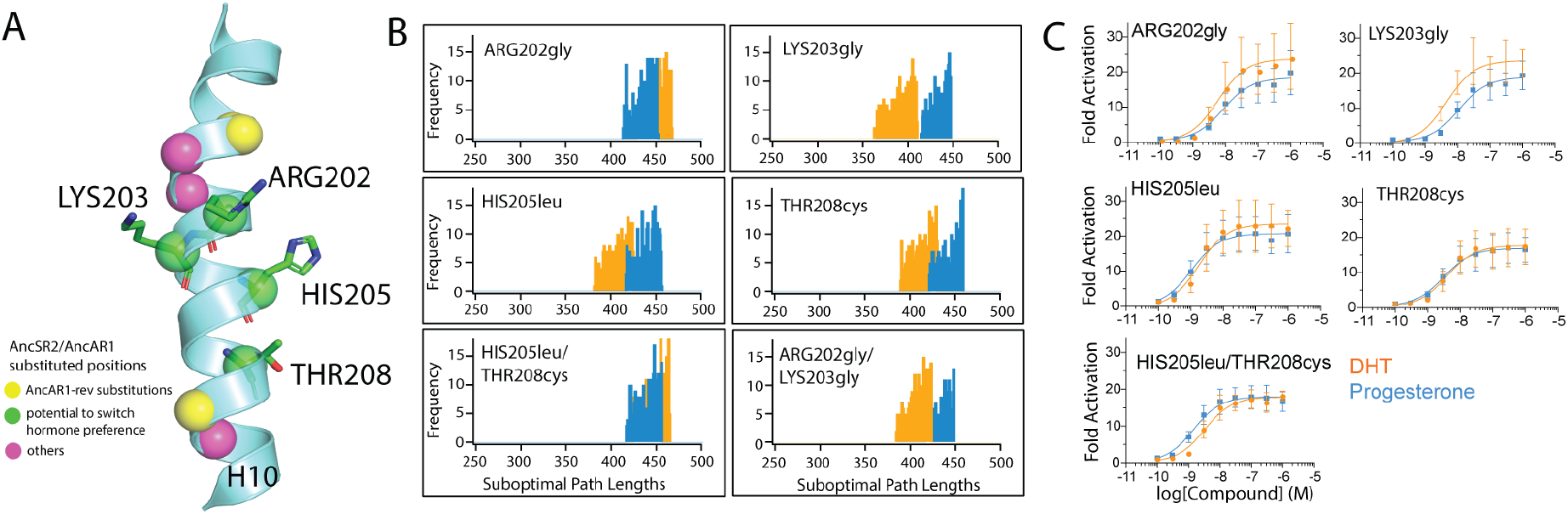
Additional H10 substitutions restore progesterone preference. A) Helix 10 residues showing AncAR1-Rev substitutions (yellow), potential preference-switching substitutions (green) and all other AncSR2-AncAR1 H10 substituted positions (magenta). B) Suboptimal paths analysis comparing strength of signaling between the ligand binding pocket and H12 for DHT and progesterone in AncAR1-Rev mutants. C) Luciferase reporter data for AncAR1-Rev mutants. The HIS205leu/THR208cys double mutant is the only one that achieves higher progesterone activation. Data shown as mean ± SEM from two biological replicates. EC_50_ values are listed in Table S3.

We next evaluated transactivation of five mutant AncAR1-Rev states: individual substitutions of the four candidates and a HIS205leu/THR208cys double-mutant of AncAR1-Rev (**Figure 6C**). Individual mutations did not demonstrate increased specific for progesterone (**Figure 6C**). However, consistent with predictions from MD simulations, the HIS205leu/THR208cys double mutant demonstrates higher specificity for progesterone than DHT (**Figure 6C**).

By analogy, reviewing the structural effects of corresponding cancer mutations at HIS874 and THR877 in human AR should reveal mechanisms by which progesterone activation is enhanced in the HIS205leu/THR208cys AncAR1-Rev mutant. Previously identified as diagnostic for androgen binding and activation, the mutation of T877 to alanine is associated with reduced androgen binding and promiscuous activation by non-androgenic hormones, including progestogens [16]. The T877C mutation in human AR was shown to broaden ligand response by increasing the size off the pocket, facilitating the entry of hormones with bulkier D-ring substituents [25]. Similar to T877, prior work reports that H874 mutations induce promiscuous ligand activation in the androgen receptor, possibly by disrupting local structure as well as by enlarging the biding pocket [26, 27].

Residue contact analysis in the MD simulations reveals that HIS205leu and THR208cys perturb a network of interactions that are crucial for communication between the ligand binding pocket and AF-2 (**Figure 7A**). While HIS205 does not interact directly with ligand, it contacts TRP72 (W741 in AR) on H5 (**Figure 7A**). Optimally positioned for interactions with H12 (M226, V234) and H3 (L43), TRP72 has also been identified as a key regulator of the shape and size of the AR binding pocket [18, 28]. The HIS205leu substitution alters the contact between position 205 and TRP72, subsequently impacting the binding pocket interaction network.

**Figure 7.**
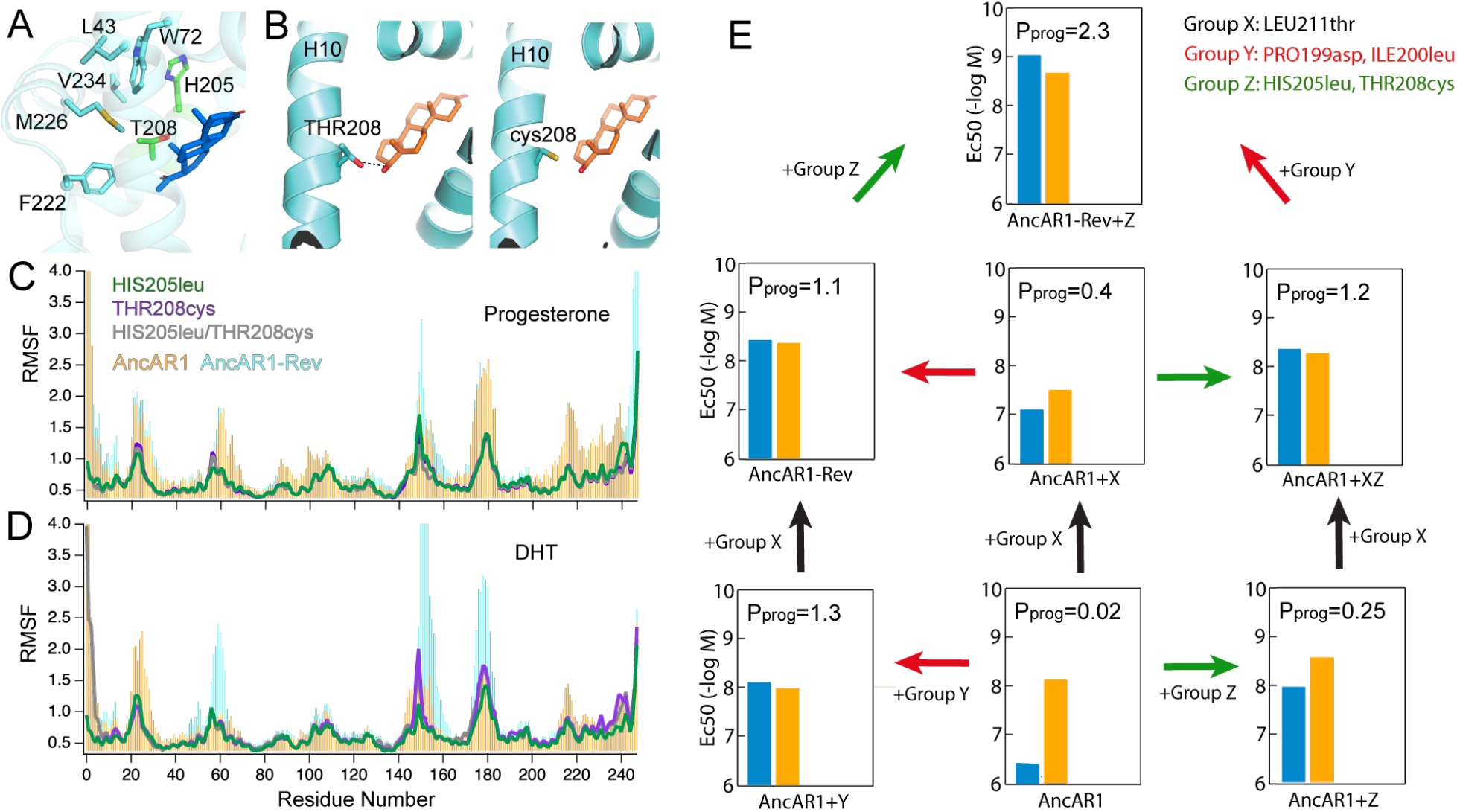
Mechanisms of restored progesterone selectivity. A) HIS205 and THR208 substitutions are positioned via W72 of H5 to interact with crucial residues on H3 (W72), H12 (M226, V234) and pre-H12 loop (F222). B) THR208cys substitution eliminates a hydrogen bond with DHT. C-D) in both progesterone and DHT complexes, individual HIS205leu (green line), THR208cys (purple line), and double (grey) substitutions lead to overall reduction in RMSF across the receptor, compared to wild type AncAR1 (orange) and AncAR1-Rev (cyan). E) Evolution of P_prog_ through sequence space. P_prog_ is calculated as DHT EC50/Prog EC50. Impact of Group X, Y and Z substitutions on progesterone (blue) and DHT (orange) activation are shown. The highest P_prog_ is only achieved when XYZ substitutions are introduced.

Conversely, THR208 contacts DHT via a C17-OH hydrogen bond, which MD simulations predict is lost upon mutation to cysteine (**Figure 7B**). Even with the loss of the hydrogen bond, cys208 remains close enough to engage the ligand in a hydrophobic interaction which is less discriminatory between DHT and progesterone. The hormone also mediates contact between H10 and the pre-H12 loop (F222) (**Figure 7A**). Simulations predict that the HIS205leu and THR208cys substitutions, individually and in combination, stabilize AncAR1 throughout the receptor, including H3, and from H10 through to the H12 (**Figure 7C, D**). Therefore, through multiple mechanisms, HIS205leu and THR208cys stabilize AncAR1-Rev while achieving a slight preference for progesterone.

To further explore the reverse evolutionary trajectory that restores progesterone preference in the HIS205leu/THR208cys AncAR1-Rev mutant, we evaluated hormone preferences in additional intermediate AncAR1 states (**Figure 7E**). We classified substitutions into three groups based on the order of their discovery: Group X (LEU211thr), Group Y (PRO199asp+ILE200leu) and Group Z(HIS205leu+THR208leu). We calculate preference for progesterone (P_prog_) in receptors using the formula P_prog_ = (DHT EC_50_/Prog EC_50_). This analysis reveals that group Z substitutions alone in the AncAR1 background are not sufficient to achieve progesterone preference. With all pairs of combinations, the highest achievable P_prog_ is 1.3, indicative of approximately equal preference for both hormones. Only when all three Groups are included does P_prog_ increase above 1. Thus, the five H10 substitutions identified here are all necessary to restore progesterone preference.

## Conclusions

The androgen receptor first appeared in the vertebrate lineage 450 million years ago in Chondrichthyes (or cartilaginous fish), the most early branching group of jawed vertebrates. AR is activated by DHT and testosterone in mammals and 11-ketotestosterone in teleost fishes [29]. All three are 17-hydroxyl steroids capable of activating shark AR, one of the earliest ARs, in luciferase assays [30]. ARs emerged from a gene duplication of the PR/AR ancestor (sometimes referred to as *AncSR3*) after androgens were already known to be present. Thus, the AR-androgen partnership evolved by molecular exploitation of an existing biomolecule [13]. However, as AncSR3 is uncharacterized, it is not known whether AR evolved via an intermediate that was strongly activated by progestogens and androgens, or whether AncAR1 represents the first androgen-specific receptor.

Previous work has characterized the mechanistic underpinning of the evolutionary shift between AncSR1 and AncSR2, the oldest two members of the SR family, as well as the evolutionary trajectories that resulted in the modern glucocorticoid [6] and mineralocorticoid receptors [8]. The model that has emerged from prior investigations into SR evolution describes two types of effects in evolutionary mutations: large effect substitutions that drive the evolution of novel ligand specificity, and small-effect substitutions fine-tune. In exploring the evolution of ligand specificity in the AR/PR clade, we find that our results are consistent with this prior description, supporting the notion of strongly conserved evolutionary mechanisms in the SR family. By examining a reversed evolutionary trajectory, we identify the LEU880thr substitution in AncAR1 as the largest effect mutation, simultaneously weakening DHT response and strengthening progesterone response.

In addition to the large effect, function-switching mutation, we identified two smaller-effect mutations that serve to restore progestogen activation to AncAR1. PRO199asp and ILE200leu fine tune ligand preference via two mechanisms: first, stabilization of the active receptor state, impacting both ligands, and second, modulating the dynamics of H10. Therefore, functional evolution in AR occurs via both ligand-specific and ligand-independent mechanisms. An attempt to study the forward effect of the thr211LEU substitution in the AncSR2 background proves futile as the large effect substitution confers constitutive activation. Similar effects have been reported in previous SR studies, allowing us to conclude that other permissive substitutions must have preceded thr211LEU in order to permit the large functional effect. Thus, this work highlights the benefit of a reverse evolutionary approach to describe trajectories for which forward studies might be limited. However, we can propose the following forward trajectory from our findings: i) AncSR2 underwent an accumulation of necessary permissive substitutions to achieve the appropriate background to tolerate functional switching; ii) H10 substitutions asp199PRO, leu200ILE, leu205HIS, and cys208THR destabilized AncSR2 for all ligands, iii) thr211LEU substitution introduces DHT-specific contacts that support activation. This proposed mechanism is consistent with a previous identification of position 211 in SRs as a D-ring sensor in the ligand binding pocket. While other SRs retain the ancestral thr211, androgen receptors are the only clade that evolved Leucine in its place, an evolutionary feature that fittingly partners the 17beta-hydroxy-D-ring-substituted androgens with a bulky, hydrophobic residue for stabilization.

However, our data could also support a revised order of events in the above proposed forward trajectory with (ii) preceding (iii), as we observe progesterone response even with leu880. Therefore, one could conceive of a model where a large effect mutation takes place after the proper background is in place, which strengthens DHT activation while maintaining modest progesterone activation (i.e., step I followed by step iii). The remainder substitutions then finetune this response, selectively decreasing progesterone activation relative to DHT (i.e., step ii). Confirming the specific sequence of evolutionary events would require an exploration of the progesterone receptor branch of the AR/PR clade, including but not limited to a resurrection of the ancestral progesterone receptor, which is beyond the scope of this study.

Perhaps the most striking result in our study is finding that all of the evolutionary, function-switching mutations occurred on a single helix, H10. Specifically, substituting various combinations of the H10 positions reported here generate AncAR1 variants with unique progesterone vs DHT responses. While the importance of H10 has been demonstrated in extant androgen receptors [31] and other steroid receptors [32, 33], this work highlights a crucial evolutionary role for this helix in both driving and finetuning the selectivity for androgens over progestogens in AR.

## Materials and Methods

### Reagents

Chemicals were purchased from Sigma Aldrich (St. Louis, MO) or Fisher (Hampton, NH). DHT was purchased from Toronto Research Chemicals (Toronto, CA). Progesterone was purchased from Sigma Aldrich. The vector for His tagged TEV was a gift from David Waugh (National Cancer Institute). The pLIC_MBP vector was a gift from John Sondek (UNC, Chapel Hill). The ancAR1 ligand binding domain (LBD) was resurrected as part of a steroid receptor ancestral reconstruction [34] and was kindly provided by Dr. Joseph Thornton (University of Chicago). The peptide corresponding to the nuclear receptor coactivator box 3 from human Tif2 was synthesized by RS Synthesis (Louisville, KY).

### Protein Expression & Purification

Protein expression and purification were performed as described in [17].

### Crystallization & Data Collection

The AR1-DHT-Tif2 ternary complex was crystallized as previously described [17]. Coordinates and structure factors were deposited in the Protein Data Bank (PDB) with the accession number 7RAE.

The AR1-Rev-Prog-Tif2 complex was crystallized as follows: prior to crystallization, an additional 500 μM Progesterone was added to the ancAR1-progesterone complex to ensure full occupancy of progesterone in the ligand binding pocket. Additionally, 500 μM of a peptide derived from human Tif2, corresponding to the nuclear receptor coactivator box 3 (740-KENALLRYLLDKDD-753), was added to the receptor-ligand complex for crystallization; yielding a 5:1 molar ratio of peptide:ancAR1. Crystals were obtained by hanging-drop diffusion in 25% PEG8000 and 50mM Tris pH 8.5. Crystals were cryoprotected in crystallant containing 30 % glycerol and were flash-cooled in liquid N_2_ at 100 K. Data to 1.55 Å were collected at the South East Regional Collaborative Access Team (SER-CAT) 21-ID at the Advanced Photon Source (APS) at Argonne National Laboratory in Chicago, Illinois, USA using a wavelength of 0.97 Å. Data were processed and scaled using HKL-2000 (HKL Research, Inc., Charlottesville, VA)[35] and phased by molecular replacement using Phaser-MR (Phenix, Berkeley, CA) (Adams et al., 2010). Data processing revealed that crystals of the ancAR1-Rev-prog-Tif2 grew in space group P4_3_2_1_2. Initial phases for the AncAR1-Rev-progesterone-Tif2 complex were determined using AncAR1-DHT-Tif2 (7RAE) as the initial search in Phenix [36]. Structure was refined using standard methods in the CCP4 suite of programs and COOT v0.9 was used for model building (MRC Laboratory of Molecular Biology, Cambridge, UK) [37]. AncAR1-Rev-prog-Tif2 coordinates and structure factors were deposited in the Protein Data Bank (PDB) with the accession number 7RAF.

### Cell Culture

U2OS cells were cultured in phenol red-free DMEM + 10 % fetal bovine serum and cultured under standard conditions (5 % CO_2_, 37 °C).

### Reporter Assays

U2OS cells were seeded at 8,000-10,000 cells per well in white-walled, clear bottom 96-well plates in phenol red-free MEMα + 10 % fetal bovine serum – Charcoal/Dextran Treated (Corning; Atlanta Biologicals). At 70-90 % confluence, cells were transfected with a construct encoding the LBD of the protein fused to the Gal4 DBD (5 ng/ well), a 9xUAS reporter construct (50 ng/ well), and a *Renilla* luciferase reporter with a CMV promoter (1 ng/ well). Transfection was performed using FuGENE at a ratio of 3:1 (FuGENE:DNA). Approximately 24 hours after transfection, compounds were dissolved in Opti-MEM and then introduced to cells to give final concentrations indicated in figures with DMSO at a final concentration of 0.370%. After ~ 24 hours, luciferase signal was quantified using the DualGlo kit (Promega). Each experiment was performed in triplicate with two biological replicates. Firefly luciferase signal was first corrected by dividing *Renilla* signal intensity for each well and then normalized relative to the DMSO control. Data was analyzed with GraphPad Prism (version 9) using a stimulating dose-response curve (three parameters – Hill slope = 1).

### Phylogenetics and ancestral sequence reconstruction

Ancestral sequence reconstruction was performed as described in [34]. Phylogenetic methods specific to AncAR1and all relevant supporting data will be added in version 2 of this preprint.

### Molecular dynamics simulations

#### Model preparation

Nineteen systems were prepared for molecular dynamics simulations using the crystal structures of the ligand binding domains of AncAR1-DHT [17] and AncAR1-rev-Progesterone, and AncSR2-progesterone (PDB 4LTW) as starting structures. The Tif2 peptide was excluded from all complexes for MD. The following AncAR1 variants were generated for simulations: a) AncAR1; b) LEU211thr-AncAR1; c) ILE200leu-AncAR1; d) PRO199asp-AncAR1; e) ILE200leu/PRO199asp-AncAR1; f) ILE200leu/LEU211thr-AncAR1, g) LEU211thr/PRO199asp-AncAR1, h) AncAR1-rev, i) HIS205leu-AncAR1-Rev, j) ARG202gly-AncAR1-Rev, k) LYS203gly-AncAR1-Rev, l) THR208cys-AncAR1-Rev, m) ARG202gly/LYS203gly-AncAR1-Rev, n) HIS205leu/THR208cys-AncAR1-Rev, o) ARG202gly/LYS203gly/ HIS205leu/THR208cys-AncAR1-Rev, p) AncSR2, q) thr211LEU-AncSR2, r) asp199PRO-AncSR2, s) leu200ILE-AncSR2. Variants were generated by manually introducing *in silico* mutations and relaxing the structure to remove unfavorable contacts. Progesterone and DHT complexes were created for each of these. The complexes were solvated in an octahedral box of TIP3P water with a 10-Å buffer around the protein complex. Na+ and Cl-ions were added to neutralize the protein and achieve physiological conditions. All systems were set up using xleap in AmberTools [38] with the parm99-bsc0 forcefield [39]. Parameters for DHT and progesterone were obtained using Antechamber [40] in AmberTools. Minimizations and simulations were performed with Amber[41] with GPU acceleration [42, 43]. Systems were minimized with 5000 steps of steepest decent followed by 5000 steps of conjugate gradient minimization with 500 kcal/mol·Å^2^ restraints on all atoms. Restraints were removed from all atoms excluding ligand atoms and the previous minimization was repeated. The systems were heated from 0 to 300 K using a 100-ps run with constant volume periodic boundaries and 5 kcal/mol·Å^2^ restraints on all protein and ligand atoms. 12 ns of MD equilibration was performed with 10 kcal/mol·Å^2^ restraints on protein and ligand atoms using the NPT ensemble. Restraints were removed and 1000 ns production simulations were performed for each system in the NPT ensemble. A 2-fs timestep was used and all bonds between heavy atoms and hydrogens were fixed with the SHAKE algorithm [44]. A cut-off distance of 10 Å was used to evaluate long-range electrostatics with Particle Mesh Ewald (PME) and for van der Waals forces. The ‘strip’ and ‘trajout’ commands of the CPPTRAJ module [45] were used to remove solvent atoms and extract 50,000 evenly spaced frames from each simulation for analysis. Root mean square fluctuations (RMSF) analysis was performed Calpha atoms of protein residues, computed for each frame in the trajectory relative to the initial structure.

### Accelerated MD

We used accelerated MD (aMD) [46] to broadly sample the conformational space in various AncAR1 complexes. This approach modifies the energy landscape by boosting the potential energy function when the true potential energy V(r) falls below a chosen level [46]. The potential function can be modified by applying a boost to the dihedral torsion or to the entire potential. We applied a dual-boosting approach that has been successful in similar studies [47]. We selected the parameters for potential energy threshold (E_P_), dihedral energy threshold (E_D_), dihedral energy boost (α_D_) and total potential energy boost (α_p_) using published guidelines [47]. All aMD calculations and simulations were performed using Amber18 [48]. Five hundred ns simulations were performed for each complex. Average dihedral energy (Eavg_D_) and average total potential energy (Eavg_p_) were obtained from classical MD simulations.

α_D_: 0.2 * (Eavg_D_ + 3.5 (kcal/mol *N_sr_)
E_D_: Eavg_D_ + 3.5 kcal/mol * N_sr_
α_p_: 0.16 kcal/mol * N_atom_
E_P_: Eavg_p_ + (0.16 kcal/mol*N_atom_) (where N_sr_ = number of total solute residues, N_atom_ = total number of atoms)

Accelerated MD was performed on 15 complexes: AncAR1, LEU211thr-AncAR1, ILE200leu-AncAR1, PRO199asp-AncAR1 and AncAR1-rev, each in complex with progesterone, DHT and in an unliganded (apo) form.

### Clustering and conformational analysis

The MMTSB toolset was used to perform clustering analyses [49] on aMD trajectories. 25,000 evenly spaced conformations were extracted from each 500 ns trajectory for clustering. Clustering was performed with a 2.6 Å cutoff. All PDBs were RMS-fit to a reference structure. Conformations from AncAR1-Rev and each single mutant AncAR1 complex (i.e. LEU211thr, ILE200leu, PRO199asp) were combined with those from the corresponding wildtype AncAR1 complex, and joint clustering was performed to identify conformations that overlap between wildtype and mutant forms of AncAR1.

#### Contact maps and Network Analysis

The NetworkView plugin in VMD [50] and the Carma program [51] were used to analyze contacts and produce dynamic networks for each system. Residue contact maps were used to determine how dynamic contacts are altered across various AncAR1 complexes [52]. To generate contact maps, all solvent atoms were stripped, leaving ligand and protein atoms. Protein residues are defined as nodes and edges (or contacts) are created between two non-neighboring nodes if the heavy atoms of the two residues are within 4.5 Å of each other for 75% of the trajectory. To produce dynamic networks, edges in residue contact maps were weighted by covariance calculated from MD simulations (following the protocol described in [53]). Edge weights inversely proportional to the calculated pairwise correlation between the nodes. The protocol described in [53] was followed.

### Suboptimal Paths

Communication between the hormone and AF-2 surface was described by generating suboptimal paths between these sites using the Floyd-Warshall algorithm [54]. Communication paths are drawn as a chain of edges connecting the ligand (source node) with a ‘sink’ node on helix 12 (E228). Due to the inverse correlation between correlation and edge weights, the sum of edges along a path between two distant nodes becomes lower as the strength of communication (i.e. correlation) increases. The optimal path is defined as the path for which the sum of edges is the lowest. However, a set of the shortest suboptimal paths, along with the optimal path, are thought to convey the greatest amount of communication between two distant nodes.

### Accession Numbers

The atomic coordinates and structure factors have been deposited in the Protein Data Bank with the accession numbers 7RAE and 7RAF for AncAR1-DHT and AncAR1-Rev-Prog complexes, respectively.

## Acknowledgements

X-ray data were collected at Southeast Regional Collaborative Access Team (SER-CAT) 22-ID beamline at the Advanced Photon Source, Argonne National Laboratory. Supporting institutions may be found at www.ser-cat.org/members/html. Use of the Advanced Photon source was supported by the U.S. Department of Energy, Office of Science, Office of Basic Energy Sciences, under Contract No. W-31-109-Eng-38.

