## Supporting Information for "Evolutionary tuning of a key helix drove androgen selectivity"

Table S1

| | AncAR1-DHT-Tif2 | AncAR1-Rev-progesterone-Tif2 |
|---|---|---|
| Resolution (highest shell) | 2.1 Å (29.41 – 2.1 Å) | 1.55 Å (32.1–1.55 Å) |
| Space Group | $P4_32_12$ | $P4_32_12$ |
| Unit Cell Dimensions | a=68.87, b=68.87, c=147.28 $\alpha = \beta = \gamma = 90$ | a=69.1, b=69.1, c=145.26 $\alpha = \beta = \gamma = 90$ |
| No. of Reflections | 21499 | 51728 |
| Rpim (highest shell) | 6.3 % (43.7 %) | 1.1 % (36.2 %) |
| $CC_{1/2}$ | n.a.[1] | 99.9 % (63.6 %)[2] |
| Completeness (highest shell) | 99.9 % (100 %) | 99.8 % (100 %) |
| Ave. Redundancy (highest shell) | 8.0 (7.3) | 7.2 (3.8) |
| $I/\sigma$ | 273.4 (18.6) | 32.8 (1.4) |
| Monomers per asymmetric unit (AU) | 1 | 1 |
| No. of protein atoms/AU | 2151 | 2132 |
| No. of ligand atoms/AU | 27 | 22 |
| No. of waters/AU | 66 | 258 |
| $R^{b}_{working}$ ($R^{c}_{free}$) | 19.3 (21.8) | 18.8 (18.9) |
| Ave. B-factors, $Å^2$ | | |
| Protein | 35.40 | 22.39 |
| Ligand | 19.65 | 27.7 |
| Water | 40.88 | 38.67 |
| r.m.s. deviations | | |
| Bond lengths, Å | 0.003 | 0.017 |
| Bond angles, ° | 0.584 | 1.47 |

[1] data collected using HKL2000_v712-Mac
[2] data collected using HKL2000_v721-Mac

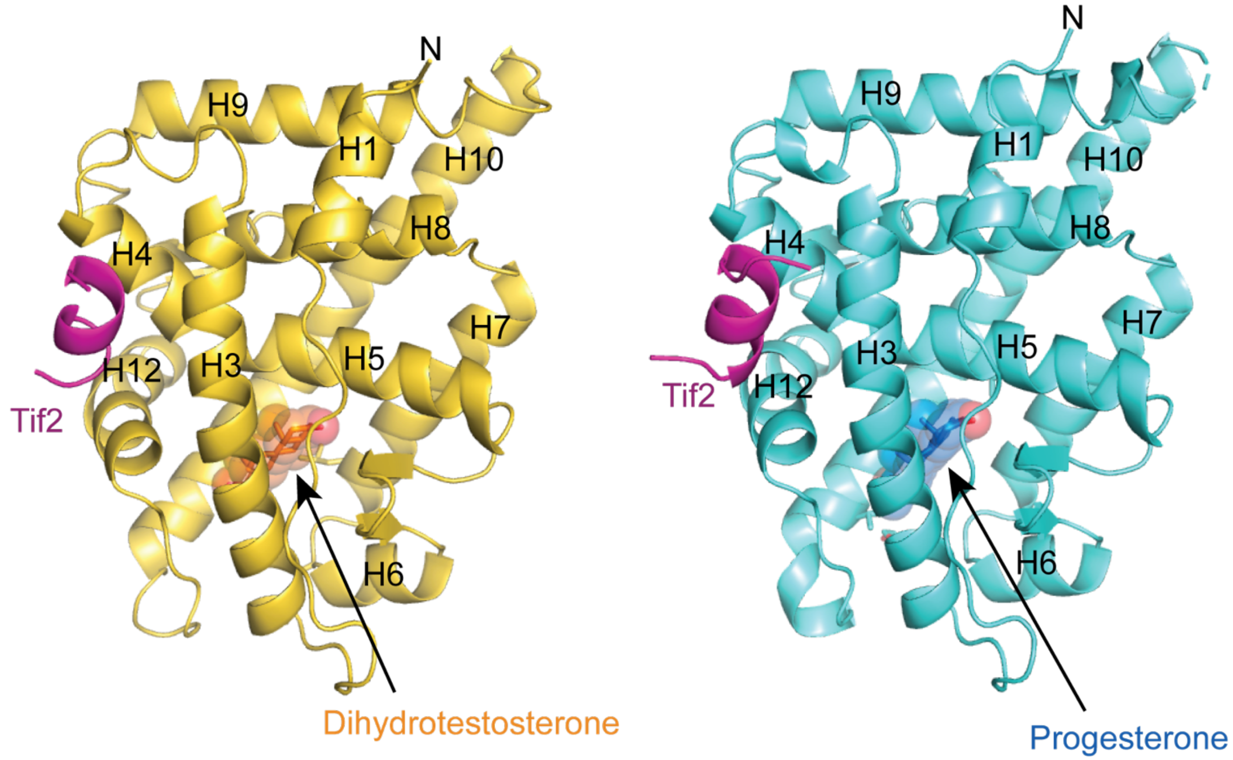


Figure S1: Structures of AncAR1-DHT-Tif2 (left) and AncAR1-Rev-Progesterone-Tif2 complexes (right). Ligands DHT (orange) and progesterone (blue) are shown in the respective binding pockets. Tif2 peptide fragment (magenta) is shown at the AF-2 surface, comprised of H12, H4 and H3.

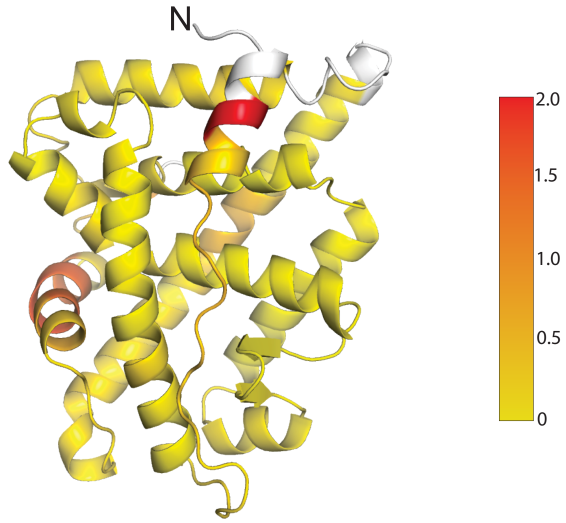


Figure S2: ProSMART analysis of pairwise comparisons of AncAR1-DHT and AncAR1-Rev-progesterone structures. Areas shown in white were not used in the comparison. Structures are colored by Procrustes score of the central residue of an aligned fragment pair according to the legend shown on the right.

| Helices | WT AR1 | | I869I | | L880t, P868d, I869I | |
|---|---|---|---|---|---|---|
| | Area interface ($Å^2$) | ΔG (kcal/mol) | Area interface ($Å^2$) | ΔG (kcal/mol) | Area interface ($Å^2$) | ΔG (kcal/mol) |
| H10 – H7 | 309.8 | -6.0 | 317.3 | -6.2 | 317.0 | -6.2 |
| H10 – H5 | 201.4 | -5.8 | 202.6 | -5.8 | 202.6 | -5.8 |
| H10 – rest of the receptor (other than H5&H7) | 1576.6 | -26.4 | 1577.8 | -26.5 | 1578.2 | -26.6 |

Table S2. Buried Surface and predicted interaction energies between H10, H5 and H7 as calculated by PISA.

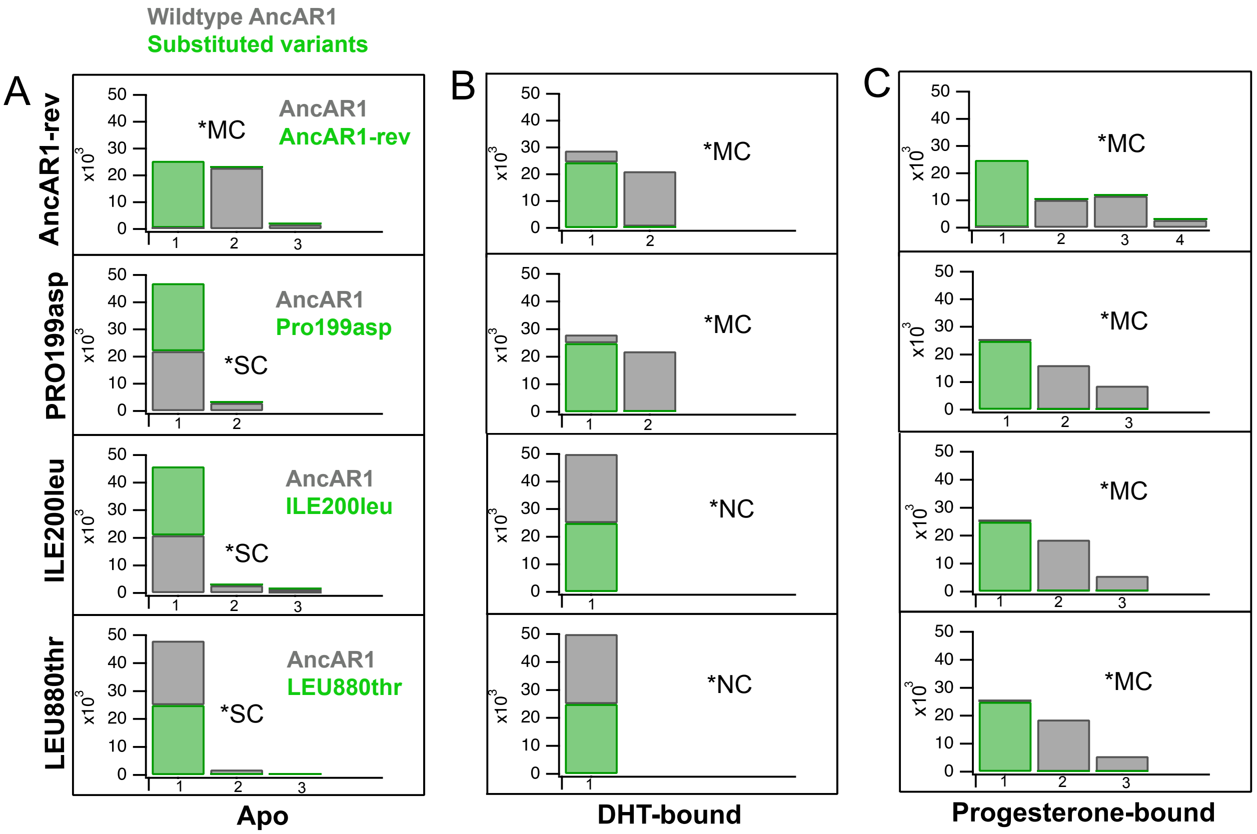


Figure S3. Conformational distributions of apo and ligand-bound AncAR1 variants. Ancestral substituted complexes (green) were co-clustered with wildtype AncAR1 (grey). Each clustering analysis is classified by the extent of overlap between wildtype and substituted AncAR1. Results are designated as Major (*MC), Small (*SC) or No (*NC) conformational changes between AncAR1 and mutants. 'Major' changes are observed when clusters are mostly 'pure', comprised entirely of either wildtype or mutant conformations/populations. 'No' changes are observed when only one mixed cluster is present, containing conformations from both wildtype and mutant AncAR1. This observation suggests that ancestral substitutions do not significantly alter conformational distributions in the receptor relative to wildtype. 'Small' changes are designated for results where the majority of conformations comprise one large mixed cluster, but a few populations fall into distinct clusters. A) Apo AncAR1 maintains a large mixed cluster in the presence of individual LEU211thr, ILE200leu or PRO1999asp substitutions, indicating that the mutations do not significantly alter its conformational distributions. All three substitutions however cause the mixed cluster to fragment into pure clusters, indicating a strong conformational and dynamic effect. B. DHT-bound AncAR1 is unaffected by Leu211thr and ILE200leu, maintaining a mixed cluster in both cases. PRO199asp and the three combined substitutions both strongly influence conformational distributions. C. Progesterone-bound DHT is affected by individual and group R substitutions, as clusters are pure in all comparisons.

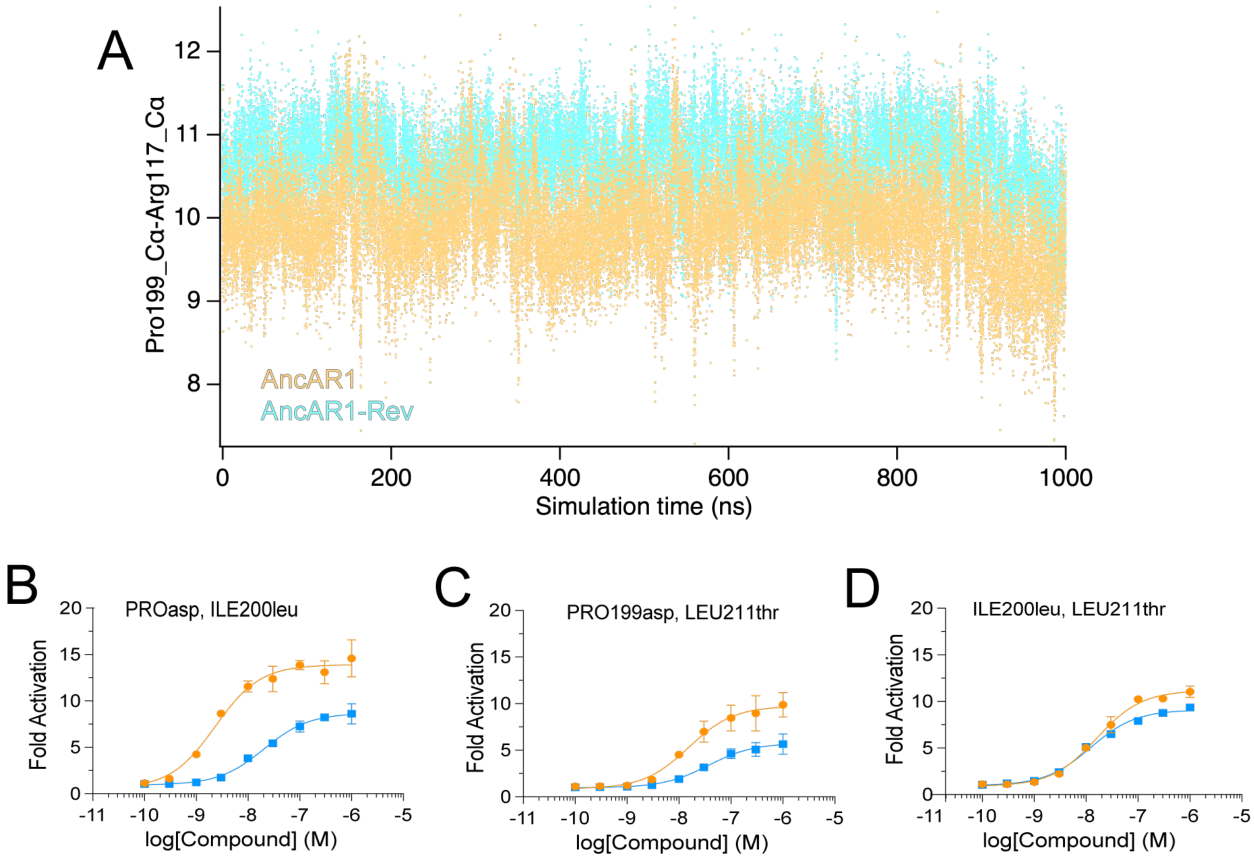


Figure S4: A) MD simulations reveal an increase in the H10-H7 interhelical distance measured between the Cα atoms of ARG117 (H7) and PRO199/asp199 (H10). B-D) Luciferase reporter activation curves for double-substituted AncAR1 mutants with DHT (orange) and progesterone (blue).

| | DHT | | Progesterone | |
|---|---|---|---|---|
| Receptor | $EC_{50}$ (nM) | Emax | $Ec_{50}$ (nM) | Emax |
| AncAR1-Rev + R202g | 5.9 (1.4, 2.7) | 23.56 | 8.5 (2.6, 29) | 17.85 |
| AncAR1-Rev + K203g | 4.6 (1.9, 11) | 23.64 | 11 (4.4, 27) | 18.43 |
| AncAR1-Rev + H205l | 1.7 (0.52, 6.1) | 23.79 | 1.1 (0.37, 3.2) | 20.77 |
| AncAR1-Rev + T208c | 3.9 (1.2, 13) | 17.51 | 3.1 (1.0, 9.4) | 16.49 |
| AncAR1-Rev + H205l+ T208c | 3.2 (1.7, 6.3) | 17.69 | 1.4 (0.66, 3.1) | 17.86 |

Table S3: EC50 values calculated from luciferase reporter assays to quantify DHT and progesterone activation of AncAR1-Rev variants.

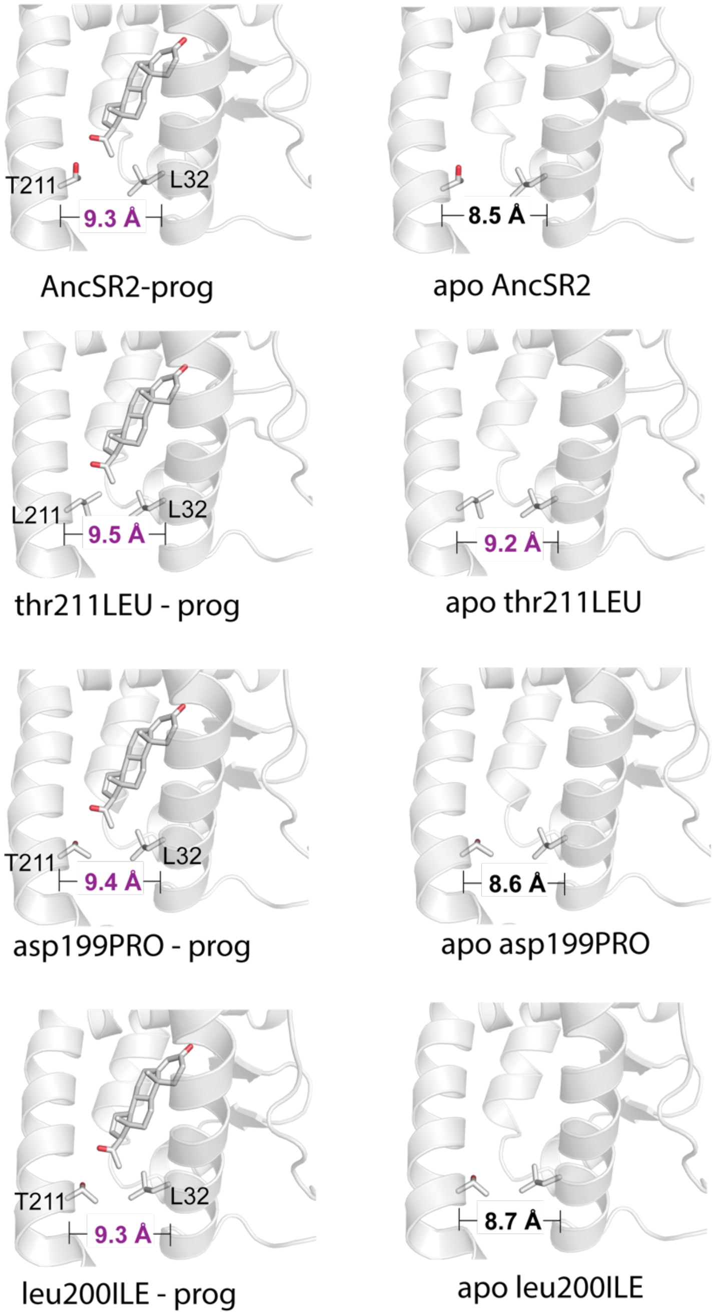


Figure S5: Changes in AncSR2 ligand binding pocket sizes upon mutation are quantified using the distance between Carbon alpha atoms of T211 (H10) and L32 (H3). Average distances calculated from MD simulations are shown for each variant. With progesterone bound, distances range from 9.3 – 9.5 angstroms for all AncSR2 variants (left). All apo complexes show a smaller pocket size (8.5 – 8.7 angstroms) except for the thr211LEU, suggesting that thr211 forms a direct contact with L32 which keeps the pocket wide in the absence of ligand.

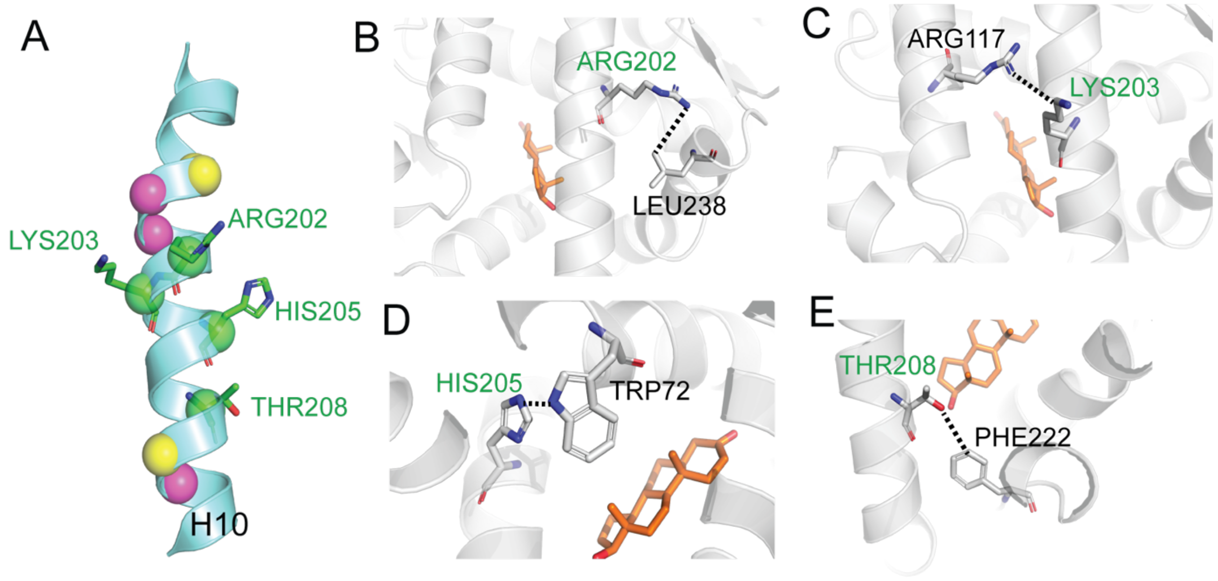


F

| | R202-L238 | | K203-R117 | | H205-W72 | | T208-F222 | |
|---|---|---|---|---|---|---|---|---|
| | DHT | Prog | DHT | Prog | DHT | Prog | DHT | Prog |
| AncAR1 | | | | | X | X | | |
| + PRO199asp | | | | | X | | | |
| + ILE200leu | X | | | | X | | X | |
| + LEU211thr | X | | | | X | | | |
| + PRO199asp + ILE200leu | X | | X | | X | | X | |
| + PRO199asp + LEU211thr | | | | | X | | X | |
| + ILE200leu + LEU211thr | X | X | | | X | | X | |
| AncAR1-Rev | X | | X | | X | X | X | |

Figure S6: To identify H10 residues which would potentially restore progesterone preference, we analyzed inter-residue contacts in our MD simulations of AncAR1, AncAR1-Rev, and all single and double substituted mutants. We compared DHT vs progesterone complexes to identify H10 residues with altered contacts. Four positions were identified (A, green), based on contacts described in B-F. B-E) For each position, the contact which displays distinct preference in DHHT vs progesterone complexes in MD simulations are shown. F) Summary of occurrences of contacts in progesterone vs DHT complexes. For example, the R202-L238 contact occurs in 5 of 8 possible DHT complexes but only one progesterone complex, suggesting that this contact might mediate hormone-specific behavior in AncAR1.
